# Wild-type U2AF1 antagonizes the splicing program characteristic of U2AF1-mutant tumors and is required for cell survival

**DOI:** 10.1101/048553

**Authors:** Dennis Liang Fei, Hayley Motowski, Rakesh Chatrikhi, Sameer Prasad, Jovian Yu, Shaojian Gao, Clara Kielkopf, Robert K. Bradley, Harold Varmus

## Abstract

We have asked how the common S34F mutation in the splicing factor U2AF1 regulates alternative splicing in lung cancer, and why wild-type U2AF1 is retained in cancers with this mutation. A human lung epithelial cell line was genetically modified so that *U2AF1*S34F is expressed from one of the two endogenous U2AF1 loci. By altering levels of mutant or wild-type U2AF1 in this cell line and by analyzing published data on human lung adenocarcinomas, we show that S34F-associated changes in alternative splicing are proportional to the ratio of S34F:wild-type gene products and not to absolute levels of either the mutant or wild-type factor. Preferential recognition of specific 3′ splice sites in S34F-expressing cells is largely explained by differential in vitro RNA-binding affinities of mutant versus wild-type U2AF1 for those same 3′ splice sites. Finally, we show that lung adenocarcinoma cell lines bearing *U2AF1* mutations do not require the mutant protein for growth *in vitro* or *in vivo*. In contrast, wild-type U2AF1 is required for survival, regardless of whether cells carry the *U2AF1*S34F allele. Our results provide mechanistic explanations of the magnitude of splicing changes observed in *U2AF1*-mutant cells and why tumors harboring *U2AF1* mutations always retain an expressed copy of the wild-type allele.

**Author Summary:** Large-scale genomics studies have identified recurrent mutations in many genes that fall outside the conventional domain of proto-oncogenes. They include genes encoding factors that mediate RNA splicing; mutations affecting four of these genes are present in up to half of proliferative myeloid disorders and in a significant number of solid tumors, including lung adenocarcinoma. Here we have characterized several properties of a common mutant version of the U2AF1 splicing factor, a component of the U2 auxiliary factor complex, in lung cells. We have found that mutant-associated changes in splice site selection are primarily influenced by the ratio of mutant and wild-type U2AF1 gene products; thus increasing wild-type U2AF1 levels represses the mutant-induced splicing program. We show that the altered splice site preferences of mutant U2AF1 can be attributed to changes in its binding to relevant 3′ splice sites. We also show that mutant U2AF1 is different from some oncogenes: the growth properties of lung cancer cell lines carrying the mutant allele are unaffected by loss of the mutant gene, while the wild-type allele is absolutely required for survival. These results advance our understanding of the molecular determinants of the mutant-associated splicing program, and they highlight previously unappreciated roles of wild-type U2AF1 in the presence of the recurrent *U2AF1*S34F mutation.

## Introduction

Somatic mutations in genes encoding four splicing factors (*U2AF1, SF3B1, SRSF2* and *ZRSR2*) have recently been reported in up to 50% of myelodysplastic syndromes (MDS) and related neoplasms and at lower frequencies in a variety of solid tumors [1–9]. Among these factors, only *U2AF1* is known to be recurrently mutated in lung adenocarcinomas (LUADs) [3, 9]. The only recurrent missense mutation of *U2AF1* in LUAD affects codon 34 and always changes the conserved serine in a zinc knuckle motif to phenylalanine (p.Ser34Phe, or S34F). This striking mutational consistency suggests a critical, yet unknown, role for *U2AF1*S34F during lung carcinogenesis. In addition, the wild-type (WT) *U2AF1* allele is always retained in cancers with common *U2AF1* mutations, including *U2AF1*S34F [2]. However, the functional significance of the wild-type allele in cells with mutant *U2AF1* is not known.

U2AF1 is a component of the U2 small nuclear ribonucleoprotein auxiliary factor complex (U2AF) [10, 11]. During early spliceosome assembly, U2AF recognizes sequences at the 3’ ends of introns to facilitate the recruitment of the U2 small nuclear ribonucleoprotein (snRNP) complex to the 3’ splice site; the recruitment occurs in conjunction with recognition of the intronic branch point by splicing factor 1 (SF1) [12, 13]. *In vitro* crosslinking assays showed that U2AF1 contacts the AG dinucleotide at the intron-exon boundary and flanking sequences [14–16].

Consistent with the critical role that U2AF1 plays in RNA splicing, *U2AF1* mutations are known to cause specific alterations in RNA splicing, most notably affecting the inclusion of cassette exons in mRNA [17–20]. However, the precise molecular basis of these splicing alterations, as well as how they are quantitatively regulated, is unknown. One possibility is that U2AF1 mutations cause altered RNA-binding affinity, resulting in altered splice site recognition. A computational model of the structure of the U2AF1:RNA complex suggested that Ser34 is a critical residue that contacts RNA [17]. Another study reported that U2AF1S34F exhibited altered affinity relative to the wild-type protein for RNA oligonucleotides derived from a cassette exon whose recognition is repressed in S34F-expressing cells [18]. Finally, the S34F mutation reportedly prevented a minimal fission yeast U2AF heterodimer from binding to a particular 3’ splice site RNA sequence [21]. However, it is not known whether altered RNA binding accounts for most S34F-associated splicing alterations and whether mechanisms other than altered binding control S34F-associated splicing.

Here, we combine genetic and biochemical approaches to show that wild-type U2AF1 antagonizes the S34F-associated splicing program in lung epithelial cells.Analyses of the transcriptomes of primary LUAD samples as well as isogenic lung cells in culture indicate that the ratio of mutant to wild-type U2AF1 gene products is a critical determinant of the magnitude of S34F-associated changes in alternative splicing. S34F-associated splicing alterations can be largely explained by differences in the relative affinities of U2AF-SF1 complexes containing mutant versus wild-type U2AF1 for RNA containing the relevant 3’ splice sites. Moreover, we show that proliferation of cancer cells with *U2AF1*S34F is critically dependent on expression of the wild-type, but not the mutant, allele of *U2AF1*.

## Results

### S34F-associated splicing correlated with S34F:WT mRNA ratios in LUAD

Before undertaking experiments to study the effects of mutant U2AF1 in cultured cells, we first examined 512 transcriptomes from primary human LUADs published by The Cancer Genome Atlas (TCGA) [9]. Thirteen of these tumors harbor the most common *U2AF1* mutation, S34F. (Two others carry rare mutations of unknown significance, S65L and G216R, and were not considered further.) We identified cassette exons whose inclusion was increased or decreased by ten percent or more in each tumor with the S34F mutation relative to the median inclusion of each cassette exon across all tumors without a *U2AF1* mutation. We then identified consensus sequence motifs for the 3’ splice sites lying immediately upstream of the cassette exons that were promoted or repressed in association with *U2AF1*S34F, represented by “sequence logos” as shown in Figs. 1A and S1. The same analysis was performed on 19 random tumors lacking a *U2AF1* mutation as controls.

**Figure 1.**
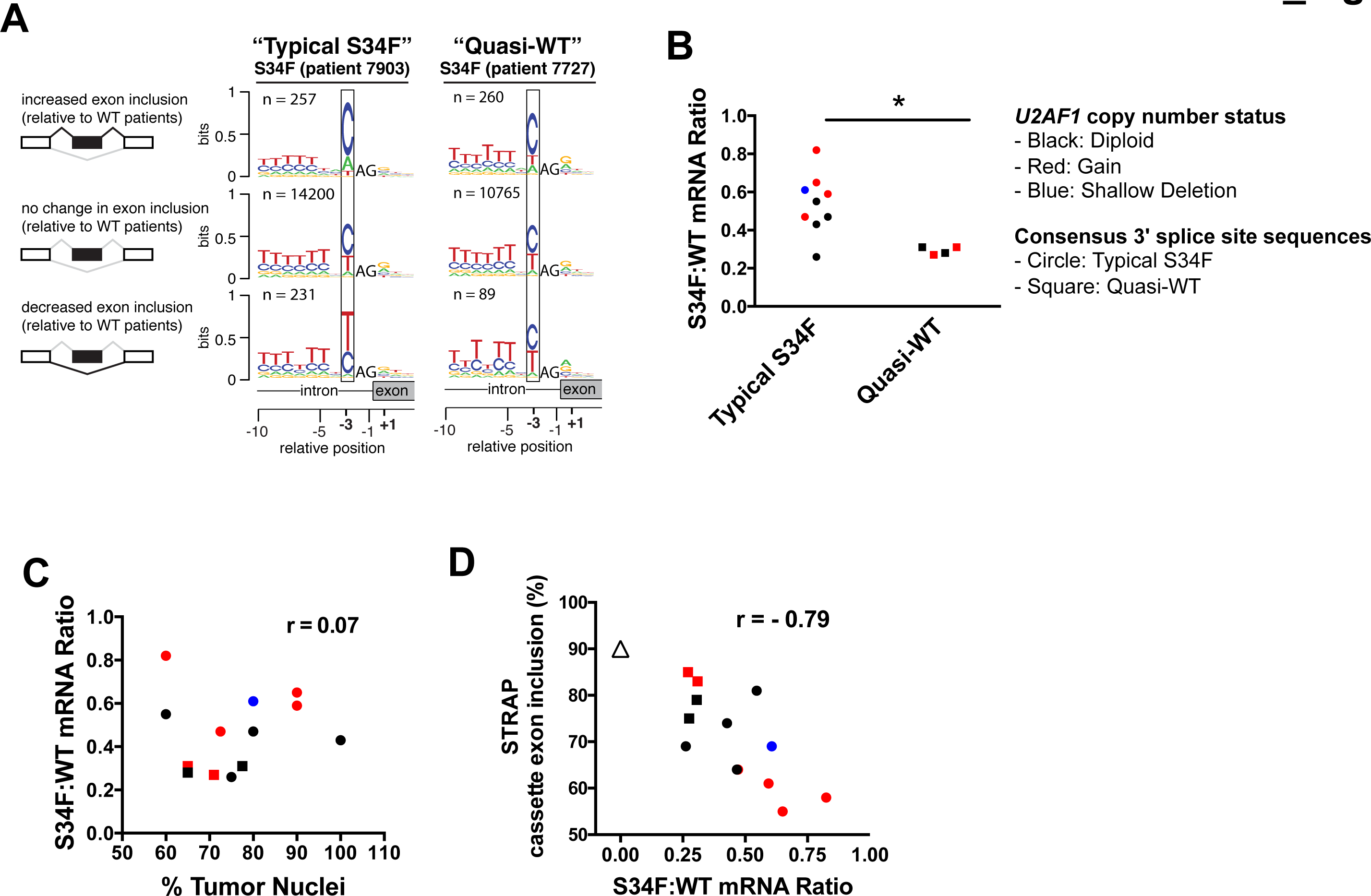
S34F-associated splicing program correlates with S34F:WT mRNA ratios in LUAD. **(A)**. Different consensus 3’ splice site sequences preceding cassette exons for two representative LUADs with the S34F mutation computed from published TCGA transcriptomes. In both cases, changes in the use of cassette exons were determined by comparisons with an average transcriptome from LUADs without *U2AF1* mutations. Boxes highlight nucleotides found preferentially at the −3 position. The nucleotide frequencies preceding exons that are more often included or more often excluded in tumors with the S34F mutation differ from the genomic consensus in the tumor from patient #7903, which represent “typical S34F” consensus 3’ splice sites, but not in the tumor from patient #7727, which represent “quasi-WT” consensus 3’ splice sites.(Consensus 3’ splice sites from transcriptomes of every S34F-mutant LUAD are presented in Supplemental Fig. S1.) The analysis was restricted to introns with 777 canonical GT-AG U2-type splice sites. The invariable AG at the 3’ splice site was not plotted to scale in order to highlight the consensus sequences at the −3 position. The vertical axis represents the information content in bits (the maximal value is two. Zero to one bit is shown). n is the number of cassette exon sequences used to construct the logo. A cartoon illustrating alternative splicing of a cassette exon (black box) is shown on the left side of the sequence logo. Black lines over splice junctions illustrate the S34F-promoted isoform for each comparison. **(B)**. *U2AF1*-mutant LUAD transcriptomes harboring “typical S34F” consensus 3’ splice sites have relatively high S34F:WT mRNA ratios. *U2AF1*-mutant LUAD samples were grouped based on the nature of the consensus 3’ splice sites. The asterisk represents a statistically significant change by Student’s t test. **(C)**. S34F:WT *U2AF1* mRNA ratios do not correlate with tumor purity in LUAD tumors with the S34F mutation. Tumor purity is represented by the percent of tumor nuclei in each LUAD sample (derived from TCGA clinical data) and plotted against the S34F:WT mRNA ratios. **(D)**. Inclusion of the *STRAP* cassette exon correlates with the S34F:WT mRNA ratio. Same as Panel **C** but the S34F:WT mRNA ratio is plotted against the inclusion frequency for the *STRAP* cassette exon. The median inclusion level of the same cassette exon for all transcriptomes from tumors without a *U2AF1* mutation (S34F:WT mRNA ratio = zero) is shown as a triangle. r, Pearson’s correlation coefficient. In panels B – D, circles represent samples with typical-S34F consensus 3’ splice site sequences; squares represent samples with quasi-WT 797 consensus 3’ splice site sequences. Colors indicate *U2AF1* copy number status as calculated by GISTIC 2.0 (See details in the Supplemental Materials and Methods): black, diploid; blue, shallow deletion; red, gain.

As illustrated by data from patient 7903 in Fig. 1A, over two hundred cassette exons were included more frequently and a similar number were included less frequently in this *U2AF1*-mutant tumor. Notably, as illustrated by the sequence logos, the nucleotide distribution at the −3 position (boxed) of promoted and repressed exons was different from that observed upstream of the much larger number of unaffected exons: A replaced T as the second most common nucleotide preceding the promoted exons, while T was more common than C in the sequence preceding the repressed exons. These patterns were observed in nine of the thirteen tumors with the *U2AF1*S34F allele (Supplemental Fig. S1A). They have been observed previously in comparisons between transcriptomes carrying the *U2AF1*S34F allele with wild-type transcriptomes [17–20], and are therefore henceforth referred to as the “typical S34F” consensus 3’ splice sites. In the other four tumors with the *U2AF1*S34F mutation, these “typical S34F” consensus 3’ splice sites were partially or completely absent (Figs. 1A,S1B). Those four tumors exhibited consensus 3’ splice sites that were similar to the 140 consensus 3’ splice sites identified in tumors lacking a *U2AF1* mutation, where variations in inclusion relative to the median wild-type sample are presumably stochastic (Supplemental Fig. S1C). Thus, we henceforth refer to these sequence patterns, as shown for tumor from patient 7727 in Fig. 1A, as “quasi-WT”.

To explain why transcriptomes of some tumors with *U2AF1* mutations showed a typical S34F-associated consensus 3’ splice sites, while others exhibited quasi-WT patterns, we estimated the levels of mutant and total *U2AF1* mRNA based on available data from the tumors to determine the ratio of mutant to wild-type (S34F:WT) mRNA.Tumors with quasi-WT patterns had low S34F:WT mRNA ratios (ranging from 0.27 to 0.31), whereas all but one tumor with the S34F-associated pattern had higher ratios (0.43 or more) (Fig. 1B). In contrast, absolute levels of *U2AF1*S34F mRNA or total *U2AF1* mRNA levels were not different between these two groups of tumors (Supplemental Figs. S2A, B).

We next sought to understand the origin of the wide range of S34F:WT ratios that we observed. These ratios, ranging from 0.26 to 0.82, could not be explained by contamination of tumor cells with non-tumor cells, since the proportion of tumor nuclei reported for these samples did not correlate with S34F:WT mRNA ratios (Fig. 1C) or with the presence or absence of the expected S34F-associated consensus 3’ splice sites (Supplemental Fig. S2C). Conversely, *U2AF1* DNA copy number correlated with the estimated levels of total *U2AF1* mRNA (Supplemental Fig. S2D). Seven of the 13 *U2AF1*S34F mutant samples, including two of the four samples displaying quasi-WT consensus 3’ splice site sequences, showed either copy number loss or gain at the *U2AF1* locus, suggesting that copy number variation (CNV) might account, at least in part, for the varying S34F:WT mRNA ratios in LUAD samples (Fig. 1B). Unbalanced allelic expression or proportions of tumor subclones might also contribute to the variable S34F:WT mRNA ratios, although these possibilities could not be readily tested using the 166 available LUAD data.

We next tested whether the *U2AF1* S34F:WT ratio was associated with quantitative changes in splicing (versus the qualitative differences in consensus 3’ splice sites described above). We correlated S34F:WT ratios with the quantitative inclusion of specific S34F-responsive cassette exons and 5’ extensions of exons resulting from competing 3’ splice sites that were reported previously [17, 19, 20]. We studied three cassette exons that exhibited less (*ASUN* and *STRAP*) or more (*ATR*) inclusion, and two 5’ extensions of exons (*FMR1* and *CASP8*) that were used less frequently in the presence of *U2AF1*S34F. Tumors with the highest S34F:WT mRNA ratios showed the lowest inclusion levels of the cassette exon in *STRAP* mRNA, whereas tumors without a *U2AF1* mutation had the highest level of inclusion (Fig. 1D; Pearson correlation of −0.79). Similar correlations were observed between inclusion levels of other representative exons and S34F:WT mRNA ratios (Supplemental Fig. S3, panels E, I, M and Q). As controls, we tested for correlations between the inclusion of these cassette exons and levels of *U2AF1*S34F mRNA, total *U2AF1* mRNA, or percent tumor nuclei.None of these analyses, with the exception of the *U2AF1*S34F mRNA level versus the inclusion of the 5’ extension of the *FMR1* exon, showed a relationship as strong as that observed for the S34F:WT mRNA ratio (Supplemental Figs. S3 panels B-D, F-H, J-L, N-P and R-T). These results indicate that the S34F:WT mRNA ratio predicts the magnitude of S34F-associated splicing in human LUAD.

### Creation of isogenic lung cell lines that recapitulated features of S34F-associated splicing in LUAD

The results presented in the preceding section, based on analyses of LUAD tumors with the *U2AF1*S34F mutation, suggest that the magnitude of S34F-associated splicing is a function of the S34F:WT mRNA ratio. To directly test this hypothesis, we developed a cell line that allows manipulation of WT and mutant U2AF1 gene product levels and measurement of the corresponding effects on RNA splicing.

The human bronchial epithelial cell line (HBEC3kt) was previously derived from normal human bronchial tissue and immortalized by introduction of expression vectors encoding human telomerase reverse transcriptase (*hTERT*) and cyclin-dependent kinase-4 (*CDK4*) [22]. To knock in a *U2AF1*S34F allele at an endogenous locus in HBEC3kt cells, we adopted a published genomic DNA editing approach [23], using the *PiggyBac* transposon that leaves no traces of exogenous DNA at the locus (Fig. 2A; see Supplemental Materials and Methods for details). We identified three cell clones at intermediate stages of gene editing after screening more than 50 primary transfectants (Supplemental Fig. S4A, B). Sanger sequencing of these intermediate clones revealed that one of the three clones carried the desired S34F missense sequence, while two clones were wild-type (Supplemental Fig. S4C). Wild-type intermediates are expected because a homologous sequence between the S34F point mutation and the drug cassette in the vector can serve as the 5’ homology arm for recombination (designated as 5’ HA#2 in Fig. 2A). From the final clones derived from these intermediates (after transposition to remove the drug cassette flanked by the *PiggyBac* elements), we chose one subclone from each of the two wild-type intermediate clones (referred to as WT1 and WT2 cells) and two subclones from the sole mutant intermediate clone (referred to as MUT1a and MUT1b cells) for all subsequent experiments with isogenic HBEC3kt cells (Supplemental Fig. S4D). The MUT and WT cells all expressed similar levels of *U2AF1* mRNA and protein (Supplemental Fig. S4E). Using high-throughput mRNA sequencing (RNA-seq) and allele-specific RT-qPCR, we observed similar levels of wild-type and mutant *U2AF1* mRNAs in the MUT cells (Figs. 2B, S5C), consistent with heterozygosity at the *U2AF1* locus.

**Figure 2.**
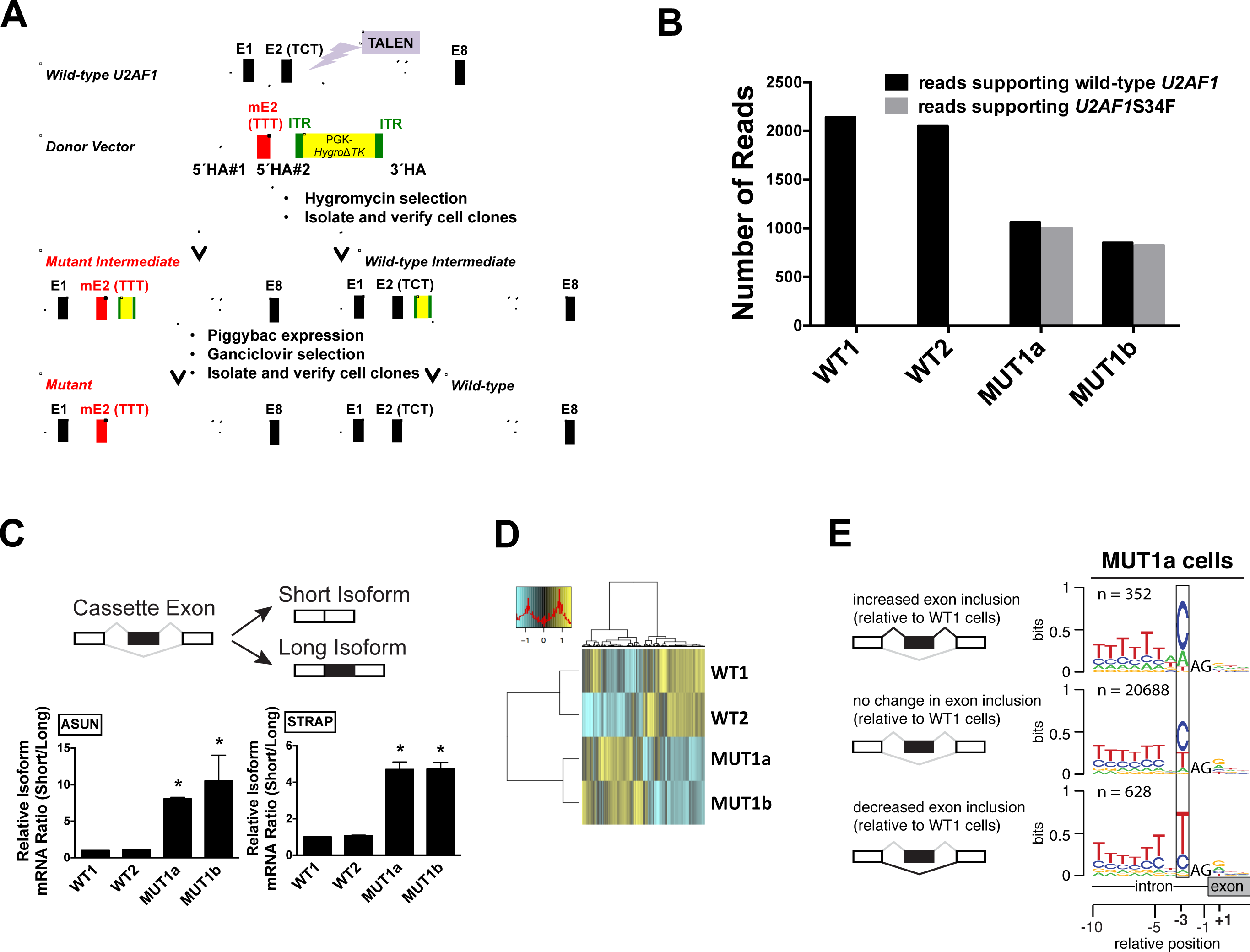
Creation of isogenic lung cell lines that recapitulate features of S34F-associated splicing. (A)Strategy to create a TCT to TTT point mutation (S34F) at the endogenous *U2AF1* locus in HBEC3kt cells. TALEN, transcription activator-like effector nuclease; E, exon; mE, mutant (S34F) exon; ITR, inverted terminal repeat; HA, homology arm. See Results and Supplemental Materials and Methods for details. **(B)**. MUT1a and MUT1b cells contain similar levels of mutant and wild-type *U2AF1* mRNA. The number of reads supporting mutant or wild-type *U2AF1* was obtained from RNA-seq, using poly(A)-selected RNA from the four cell lines. **(C).** The S34F-associated cassette exons in *ASUN* and *STRAP* mRNAs show decreased inclusion in MUT cell lines. (Top) Scheme of alternative splicing with a cassette exon (black box) to generate short and long isoforms in which the cassette exon is excluded or included. (Bottom) Alternative splicing of cassette exons in *ASUN* and *STRAP* mRNAs, measured by RT-qPCR using isoform-specific primers. The short/long isoform ratio in WT1 cells was arbitrarily set to 1 for comparison. Asterisks represent statistical significant changes as compared to that in WT1 cells. Error bars represent s.e.m. (standard error of the mean) (n = 4). **(D)**. Heat 818 map depicting the inclusion levels of all cassette exons that showed at least a 10% change in use among the cell lines. The dendrogram was constructed from an unsupervised cluster analysis. **(E).** Sequence logos from 3’ splice sites preceding cassette exons with altered inclusion in MUT1a cells display typical S34F consensus 3’ splice sites. Logos were constructed as in Fig. 1A based on the transcriptome of MUT1a cells in comparison with that of WT1 cells. Other comparisons of transcriptomes from MUT and WT cell lines yielded similar sequence logos.

To determine how the engineered *U2AF1 S34F* allele affects mRNA splicing, we first assayed the inclusion levels of 20 cassette exons that were previously reported to be associated with mutant *U2AF1* in both LUAD and AML (acute myeloid leukemia) [19]. We confirmed that all 20 of these cassette exons, which included the previously studies *ASUN* and *STRAP* cassette exons, were indeed S34F-dependent in our engineered cells using RT-qPCR with isoform-specific primers (Figs. 2C, S6).

We next evaluated the global difference in cassette exon recognition in MUT versus WT cells using RNA-seq (Supplemental Table S1). MUT and WT cells were grouped separately in an unsupervised cluster analysis based on cassette exon inclusion (Fig. 2D). We observed the expected consensus 3’ splice sites of cassette exons that were promoted or repressed in MUT versus WT cells (Fig. 2E). Overall, these results indicate that we successfully created clonal HBEC3kt cells isogenic for *U2AF1*S34F and that the MUT cells exhibited similar alterations in splicing relative to their WT counterparts as do primary LUAD transcriptomes.

### The ratio of S34F:WT gene products controlled S34F-associated splicing in isogenic lung cells

We next used our isogenic cell model with the *U2AF1*S34F mutation to test the hypothesis that the S34F:WT ratio, rather than absolute levels of the mutant or wild-type gene products, controls S34F-associated splicing. We tested two specific predictions.First, changing the levels of *U2AF1*S34F while keeping the S34F:WT ratio constant should not affect S34F-associated splicing. Second, changing the level of wild-type *U2AF1* while keeping the level of *U2AF1*S34F constant (e.g., allowing the S34F:WT ratio to change) should alter the inclusion of S34F-dependent cassette exons. We 241 tested these predictions in the isogenic HBEC3kt cells by manipulating levels of wild-type or mutant *U2AF1* gene products and measuring the subsequent changes in S34F-associated splicing.

We first reduced the amounts of both mutant and wild-type *U2AF1* mRNA concordantly in MUT1a cells, keeping the S34F:WT mRNA ratio constant. This was achieved by transducing MUT1a cells with short hairpin RNAs (shRNAs) that target regions of the *U2AF1* transcripts distant from the S34F missense mutation. The same shRNAs were also introduced in WT1 cells as a control. Allele-sensitive RT-qPCR confirmed that the mRNA ratio of the mutant and wild-type *U2AF1* remained constant (Supplemental Fig. S7A), while the overall U2AF1 mRNA and protein levels were reduced by more than 90% (Figs. 3A, bottom panel, and S7B). Knockdown of total *U2AF1* in both MUT1a and WT1 cells did not cause significant changes in recognition of the *ASUN* or *STRAP* cassette exons, two splicing events that are strongly associated with *U2AF1*S34F, in either cell line (Fig. 3A, upper panels). Similar results were obtained for two additional S34F-associated cassette exons in *USP25* and *AXL* that exhibit increased inclusion in cells expressing *U2AF1*S34F (Supplemental Fig. S7C, D).

**Figure 3.**
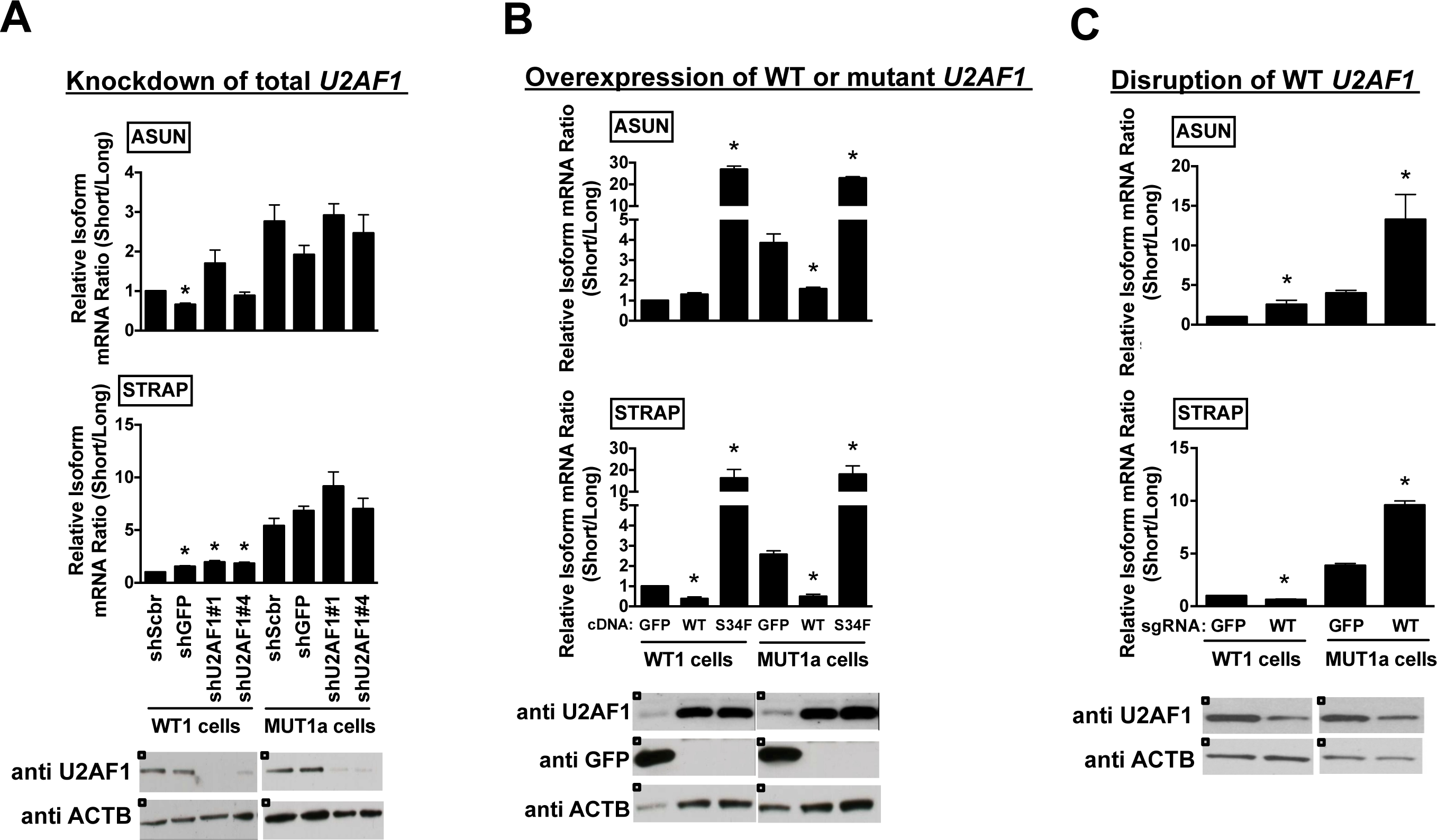
The ratio of S34F:WT U2AF1 gene products controls S34F-associated splicing in isogenic HBEC3kt cell lines. (A)Reduction of both mutant and wild-type U2AF1 RNA and protein does not affect S34F-associated splicing. WT1 and MUT1a cells were transduced with shRNAs against *U2AF1* (shU2AF1#1 and #4) or two control shRNAs (shScbr, an scrambled shRNA; shGFP, an shRNA against *GFP*). Total RNA and protein were harvested 4 days later.The frequencies of incorporation of cassette exon sequences in *ASUN* and *STRAP* mRNAs (top and middle panels) were determined by the relative short/long isoform ratios by RT-qPCR, as represented in Fig. 2C. Immunoblots for U2AF1 and ACTB in total cell lysates are shown in the bottom panel. Asterisks represent statistical significant changes as compared to shScbr-transduced condition in each cell line. Error bars represent s.e.m (n = 3). **(B)**. Overexpression of wild-type or mutant *U2AF1* to change S34F:WT ratios alters S34F-sensitive splicing. WT1 and MUT1a cells were transduced with expression vectors encoding *GFP*, wild-type (WT) or mutant (S34F) *U2AF1* for 3 days before harvesting cells to quantify the level of splicing changes and proteins as in panel A. Asterisks represent statistical significant changes as compared to GFP-transduced condition in each cell line. Error bars represent s.e.m (n = 3). **(C)**. Disruption of WT *U2AF1* by gene editing to increase S34F:WT ratios enhances S34F-sensitive splicing. WT1 and MUT1a cells were transduced with Cas9 and either sgRNA-GFP or sgRNA-WT. Total RNA and protein were harvested 6 days later for assays as in panel **A**. Asterisks represent statistical significant changes as compared to Cas9 and sgRNA-GFP-transduced condition in each cell line. Error bars represent s.e.m. (n = 3).

We next confirmed that knockdown of *U2AF1* was sufficient to alter splicing events known to be dependent on wild-type U2AF1. We studied a competing 3’ splice site event in *SLC35C2*, in which the use of an intron-proximal over an intron-distal 3’ splice site depends on the level of U2AF1 independent of a *U2AF1* mutation [24].Knockdown of total *U2AF1* in either WT1 or MUT1a cell lines reduced the use of the U2AF1-dependent intron-proximal 3’ splice site (Supplemental Fig. S7E). Thus, reduction of *U2AF1* to levels sufficient to affect U2AF1-dependent alternative splicing did not affect S34F-associated splicing in MUT1a cells when the S34F:WT ratio was maintained.

We next altered the S34F:WT ratio by separately overexpressing mutant or wild-type *U2AF1* in WT1 and MUT1a cells and examining the subsequent changes in the recognition of the *ASUN* and *STRAP* cassette exons. These cassette exons are preferentially excluded in cells expressing *U2AF1*S34F. Increasing the amount of U2AF1S34F protein—hence increasing the S34F:WT ratio in either cell type—further enhanced skipping of these cassette exons (Fig. 3B). Conversely, decreasing the S34F:WT ratio in MUT1a cells by increasing the production of wild-type U2AF1 protein reduced the extent of exon skipping to approximately the same levels seen in WT1 cells (Fig. 3B).

We also altered the production of wild-type U2AF1 mRNA and protein in MUT1a cells by disrupting the endogenous wild-type *U2AF1* locus with the CRISPR-Cas9 system. Single-guide RNAs (sgRNAs) designed to match either the WT or mutant *U2AF1* sequences were shown to disrupt either reading frame selectively, generating indels (insertions and deletions) at the *U2AF1* locus and thereby changing the S34F:WT ratios (Supplemental Figs. S8, S9). Since WT *U2AF1* is required for the growth of these cells (as shown below; Fig. 6), we extracted RNA and protein from cells six days after transduction with Cas9 and sgRNA-WT, when depletion of wild-type U2AF1 was incomplete (Fig. 3C). Selective disruption of wild-type *U2AF1* increased the S34F:WT mRNA ratio in MUT1a cells; as predicted, the extent of exon skipping was further increased in *ASUN* and *STRAP* mRNAs (Fig. 3C). Notably, the degree of exon skipping induced by mutant cDNA was similar to that caused by disrupting the wild-type *U2AF1* allele (compare Figs. 3C with 3B), even though the absolute protein levels of U2AF1S34F were different in the two experiments. These results show that wild-type U2AF1 antagonizes the activity of U2AF1S34F by a competitive mechanism, such that the S34F:WT ratio controls the magnitude of S34F-associated splicing changes independent of levels of either mutant or wild-type protein.

### Disruption of WT U2AF1 globally enhanced S34F-associated splicing

Results in the preceding sections are based on studies of a few well-documented S34F-responsive cassette exons that likely serve as surrogates for the global effects of U2AF1S34F on splice site recognition. To determine whether these results reflect general rules governing S34F-associated splicing, we used RNA-seq to measure the consequences of altering S34F:WT ratios on global recognition of cassette exons.When wild-type *U2AF1* was diminished by CRISPR-Cas9-mediated disruption in MUT1a cells (see Fig. 3C), the S34F-associated changes in inclusion (Fig. 4A) or exclusion (Fig. 4B) of cassette exons were enhanced. These global effects are consistent with our measurements of individual cassette exons by RT-qPCR (Fig. 3C). An unsupervised cluster analysis suggests that disruption of wild-type *U2AF1* in MUT1a cells primarily enhances the magnitude of changes for S34F-associated cassette exons (Fig. 4C). We also observed a modest increase in the preference for C versus T at the −3 position of the consensus 3’ splice sites of cassette exons that were promoted versus repressed in association with *U2AF1*S34F (Fig. 4D) following reduction of wild-type *U2AF1* levels, consistent with the observed association between the S34F:WT mRNA ratio and typical S34F-associated consensus 3’ splice sites identified in LUAD tumor transcriptomes (Fig. 1).

**Figure 4.**
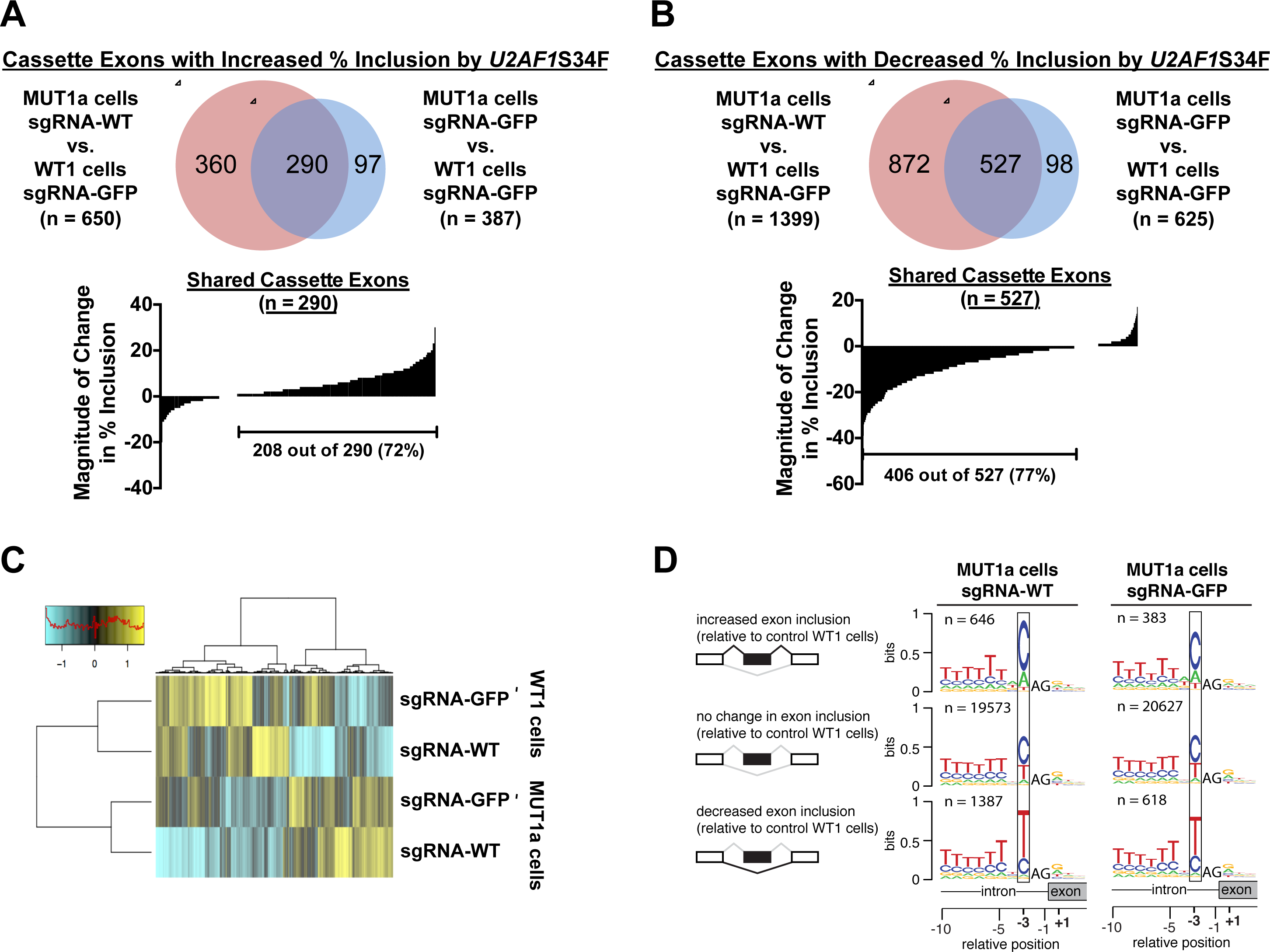
Increasing the ratio of S34F:WT gene products by disrupting the wild-type *U2AF1*locus enhances S34F-associated splicing of cassette exons. (A). RNA-seq was performed for WT1 and MUT1a cells transduced with Cas9 and either sgRNA-GFP or sgRNA-WT. Upper panel: The Venn diagrams show overlap of cassette exons that display at least a ten percent increase in inclusion levels in MUT1a cells relative to levels in WT1 cells, with or without CRISPR-Cas9-mediated disruption of wild-type *U2AF1*. Lower panel: Increasing the S34F:WT ratio by disrupting wild-type *U2AF1* enhances the magnitude of S34F-associated splicing. The “waterfall” plot depicts changes in percent inclusion levels for all shared cassette exons identified from the Venn diagram when the wild-type *U2AF1* locus was disrupted. Each vertical bar represents one shared cassette exon. **(B)**. The analysis shown in Panel A was repeated for cassette exons showing ten percent or more decreased inclusion in MUT1a cells. **(C)**. Heat map depicting the inclusion levels of all cassette exons that showed at least a 10% change in use among the treatment conditions. The dendrogram was constructed from an unsupervised cluster analysis. (D). Enhanced features of S34F-associated logos at 3’ splice acceptor sites after disruption of wild-type *U2AF1*.Sequence logos were constructed as in Fig. 1A, based on the indicated comparisons.

We observed similar results when we extended our analyses to include other types of alternative splicing beyond cassette exon recognition (Supplemental Fig. S10). We used RT-qPCR to validate five splicing alterations that are sensitive to ablation of wild-type *U2AF1* in the presence of *U2AF1*S34F (Supplemental Fig. S11). Two of the five events involve incorporation of cassette exons (in *ATR* and *MED15*), two involve competition between different 3’ splice sites (in *CASP8* and *SRP19*), and one involves a choice between two mutually exclusive exons (in *H2AFY*). We conclude that the importance of the S34F:WT ratio for S34F-dependent splicing changes extends from cassette exon recognition to other types of alternative splicing.

### Changes in RNA splicing correlated with relative binding affinities of mutant and WT U2AF1 complexes

Based on the role of U2AF1 in 3’ splice site recognition, we hypothesized that differential RNA binding by mutant and WT U2AF1 could contribute to the observed dependence of S34F-associated splicing on the WT:S34F ratios. It has previously been shown that the S34F mutation reduces the binding affinity of the U2AF1-containing complex for a representative skipped splice site [18]. However, whether changes in RNA binding could account for exon inclusion, as well as the generality of this observation, were unknown. We therefore tested whether altered RNA-binding affinity could explain mutation-dependent increases (*ZFAND1*, *FXR1, ATR, MED15*) and decrease (*CEP164*) in exon inclusion, as well as competing 3’ splice site recognition (*FMR1*). These splicing events exhibited S34F-associated alterations in both our isogenic cell lines (Supplemental Table S1) and in LUAD transcriptomes (Supplemental Table S2).

We determined the RNA binding affinities of purified U2AF1-containing protein complexes using fluorescence anisotropy assays, in which the anisotropy increases of fluorescein-labeled RNA oligonucleotides following protein titration were fit to obtain the apparent equilibrium binding affinities (Supplemental Fig. S12; Supplemental Materials and Methods). The recombinant proteins comprised either WT or S34F-mutant U2AF1 as ternary complexes with the U2AF2 and SF1 subunits that recognize the adjoining 3’splice site consensus sequences. The constructs were nearly full length and included the relevant domains for 3’ splice site recognition [15, 25, 26] (Fig. 5A). The six pairs of tested RNA oligonucleotides (33 − 35 nucleotides in length) were derived from the proximal or distal 3’ splice sites of the six genes listed above (Figs. 5B, S12). Combined with prior results for sequence variants derived from the S34F-skipped *DEK* cassette exon [18], we have in total measured binding affinities for 16 RNA oligonucleotides, which consist of five sequences with a U at the −3 position of the 3’ splice site (“UAG” splice sites), seven “CAG” splice sites, and four “AAG” splice sites.

**Figure 5.**
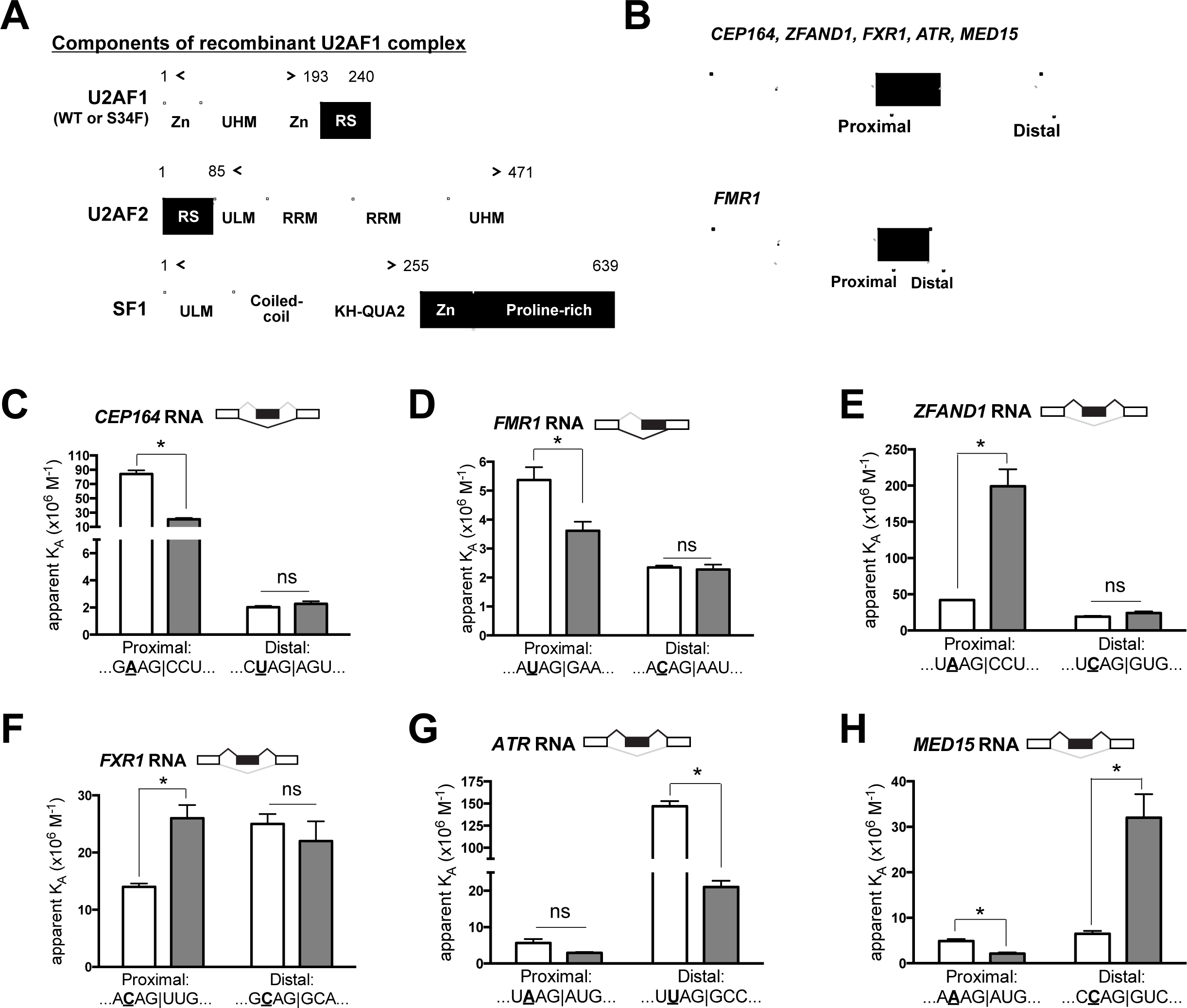
Differential binding of mutant and wild-type 2AF1 complexes to RNA oligonucleotides explains most S34F-associated alterations in RNA splicing. (A) Cartoons illustrate components of recombinant U2AF1 complexes used in the binding assay. Full-length proteins are shown but only partial sequences (denoted by bi-directional arrows) were used to make recombinant protein complexes. KH-QUA2, K-Homology Quaking motif; RRM, RNA recognition motif domain; RS, arginine/serine-rich domain; UHM, U2AF homology motif; ULM, U2AF ligand motif; ZnF, zinc finger domain. (B)Scheme of alternative splicing patterns for cassette exons (top diagram) and 5’ extended exons from competing 3’ splice site selection (bottom). The brackets indicate the positions of RNA oligonucleotides used for the binding assays. Exons are shown as boxes: white boxes indicate invariant exonic sequences and black boxes denote sequences that are incorporated into mRNA (exonic) only when the proximal 3’ splice sites are used. Introns are shown as solid lines. The grey lines represent possible splices. The names of characterized genes that conform to the patterns shown in the upper and lower cartoons are indicated. **(C − H)**. Mutant and wild-type U2AF1 complexes have different affinities (K_A_’s) for relevant 3’ splice site oligonucleotides. To accomplish the binding assays, the wild-type or mutant U2AF1 protein complexes were titrated into 5’ fluorescein-tagged RNA oligonucleotides over a range of concentrations as described in the Supplemental Materials and Methods. RNA sequences from −4 to +3 relative to the 3’ splice sites (vertical lines) in proximal and distal positions are shown. The nucleotide at the −3 position is bolded and underlined. Empty bars, K_A_ for WT U2AF1 complex; grey bar, K_A_ for mutant U2AF1 complex. For ease of comparison between the affinity binding results with S34F-associated splicing, S34F-promoted splices, as determined from RNA-seq data, are shown on top of each bar graph in black 891 lines. The fitted binding curves, full oligonucleotide sequences, and apparent equilibrium dissociation constants are shown in Supplemental Fig. S12. The relative changes in affinity and use of proximal *versus* distal splice sites are summarized in Supplemental Table S3. The asterisk represents a statistically significant change by unpaired t-tests with Welch’s correction. ns, not statistically significant.

The trends among S34F-altered RNA binding affinities of U2AF1 complexes (Fig.5C − H) for the tested splice site sequences generally agreed with the nucleotide distributions observed in consensus 3’ splice site that are promoted or repressed by *U2AF1*S34F (Figs. 1A, S1A). The S34F mutation reduced the affinities of U2AF1-containing complexes for four out of five tested “UAG” splice sites, consistent with T as the most common nucleotide at the −3 position of the 3’ splice site (henceforth referred to as −3T) for S34F-repressed exons. Likewise, the S34F mutation often increased the affinities of U2AF1 complexes for “CAG” splice sites (for three out of seven tested sequences), consistent with −3C as the most common nucleotide preceding S34F-promoted exons. The remaining tested “UAG” or “CAG” splice sites showed no significant difference between S34F and WT protein binding. The “AAG” splice sites lacked a consistent relationship to the S34F-induced RNA affinity changes of U2AF1 complexes *per se*. However, the S34F mutation increased the binding of the U2AF1-containing complex for one “AAG” splice site for an S34F-promoted exon (*ZFAND1*),which is consistent with the greater prevalence of −3A than −3T preceding S34F-promoted exons.

Overall, the altered binding affinities of U2AF1-containing complexes for the proximal 3’ splice site could account for four of the six S34F-associated alternative splicing events (*CEP164*, *FMR1*, *ZFAND1*, *FXR1*) (Figs. 5C − F, Supplementary Table S3). Similar to the previously-tested S34F-skipped splice site in *DEK* [18], the S34F mutation decreased affinities of the U2AF1-containing complexes for the skipped 3’ splice sites of the *CEP164* and *FMR1* exons (Fig. 5C, D). Remarkably, the S34F mutation enhanced binding of the U2AF1-containing complexes to the proximal 3’ splice sites of *ZFAND1* and *FXR1* (Fig. 5E, F), which could explain the enhanced inclusion of these exons in cell lines and LUAD (Supplementary Table S3). In agreement with the observed splicing changes, the S34F mutation had no significant effect on the distal 3’ splice sites of these exons.

The affinities of the mutant U2AF1 complexes for the proximal splice site oligonucleotides of the remaining two S34F-promoted exons (*ATR* and *MED15*) were either similar to wild-type or decreased (Fig. 5G, H). These results differed from the S34F-dependent increase in U2AF1 binding to the proximal 3’ splice site that was observed for *ZFAND1* and *FXR1* (Fig. 5E, F), which could readily explain the enhanced exon inclusions. However, for the *ATR* pre-mRNA, the *U2AF1* mutation reduced binding to the distal more than to the proximal 3’ splice site (Fig. 5G, third and fourth columns).Given co-transcriptional splicing in the 5’-to-3’ direction, the downstream (as opposed to upstream) 3’ splice sites could compete as splicing acceptors for a given 5’ donor splice site when transcription is relatively rapid as compared to splicing. (Such 3’ splice site competition likely occurs relatively frequently, as the *ATR* cassette exon is alternatively spliced even in wild-type cells). As such, a “net gain” in affinity for the proximal relative to distal 3’ splice site could explain the observed S34F-associated exon inclusion in *ATR* mRNAs. For the *MED15* pre-mRNA, deviation of the S34F-associated splicing changes and a simple RNA affinity model suggested that additional mechanisms control selection of the *MED15* 3’ splice sites. In total, our measurements of *in vitro* RNA-binding affinities are sufficient to explain six of seven tested alterations in splice site recognition driven by *U2AF1*S34F (Fig 5C-H and in [18]).

### HBEC3kt and LUAD cells were not dependent on *U2AF1*S34F for growth, but wild-type *U2AF1* was absolutely required

Other than its effect on RNA splicing, the consequences of the *U2AF1*S34F mutation on cell behavior are largely unknown. Recurrent mutations, such as *U2AF1 S34F*, are considered likely to confer a selective advantage to cells in which they occur when expressed at physiologically relevant levels. However, mutant HBEC3kt cells (MUT1a and MUT1b) do not exhibit obvious phenotypic properties of neoplastic transformation—such as a growth advantage over wild-type cells (Supplemental Fig.S13) or an ability to grow in an anchorage-independent manner—that are frequently observed in cultured cells expressing well-known oncogenes, like mutant *RAS* genes.

Another attribute of some well-known oncogenes, such as *BCR-ABL* fusion in chronic myeloid leukemia or mutant *EGFR* or *KRAS* in LUAD, is the dependence on sustained expression of those oncogenes for the maintenance of cell growth or viability.To determine whether LUAD cells harboring a pre-existing S34F mutation are dependent upon (or “addicted to”) the mutant allele, we searched the COSMIC database for LUAD cell lines with the *U2AF1*S34F allele [27]. Two cell lines (H441 and HCC78) were found and both these cells exhibit copy number gains at the *U2AF1* locus three copies for H441 cells; four copies for HCC78 cells) and one copy of a variant allele. We confirmed that *U2AF1*S34F was the minor allele in these cells by Sanger sequencing and allele-specific RT-qPCR (Supplemental Fig. S14A, B). We further used the CRISPR-Cas9 system to selectively disrupt the wild-type or mutant *U2AF1* sequences and then assessed the impact of inactivating the *U2AF1* alleles on the clonogenic growth of the two LUAD lines with the *U2AF1* mutation. In addition, we performed similar experiments with the LUAD cell line A549 (wild-type for U2AF1) and the HBEC3kt-derived MUT1a cell line.

In all instances, loss of the mutant allele did not impair cell growth. Only one line (H441) exhibited altered growth, in the form of a two-fold increase in clonogenicity (Fig. 6A). Successful disruption of the *U2AF1*S3F allele was confirmed by restoration of a normal RNA splicing profile in subclonal cells derived from the clonogenic assay colonies (see Supplemental Figs. S14 – S17, Tables S4 and S5, and text below). In contrast, loss of the wild-type allele completely inhibited clonogenic growth in all tested cell lines, regardless of whether the line carried the *U2AF1*S34F allele or not. A rescue experiment confirmed that loss of cell growth was due to loss of wild-type *U2AF1* expression. The loss of clonogenic capacity after disrupting endogenous *U2AF1* in A549 cells was prevented by first transducing them with a form of wild-type *U2AF1* cDNA (Fig. 6B) that is not predicted to be a target for sgRNA-WT (Supplemental Fig. S8). Overall, these findings indicate that wild-type *U2AF1* is required for the clonogenic growth of cells, including lung cancer cell lines, that the S34F mutant is unable to compensate for loss of the wild-type allele, and that LUAD cells with the S34F mutation are not dependent on the mutant allele for growth *in vitro*.

**Figure 6.**
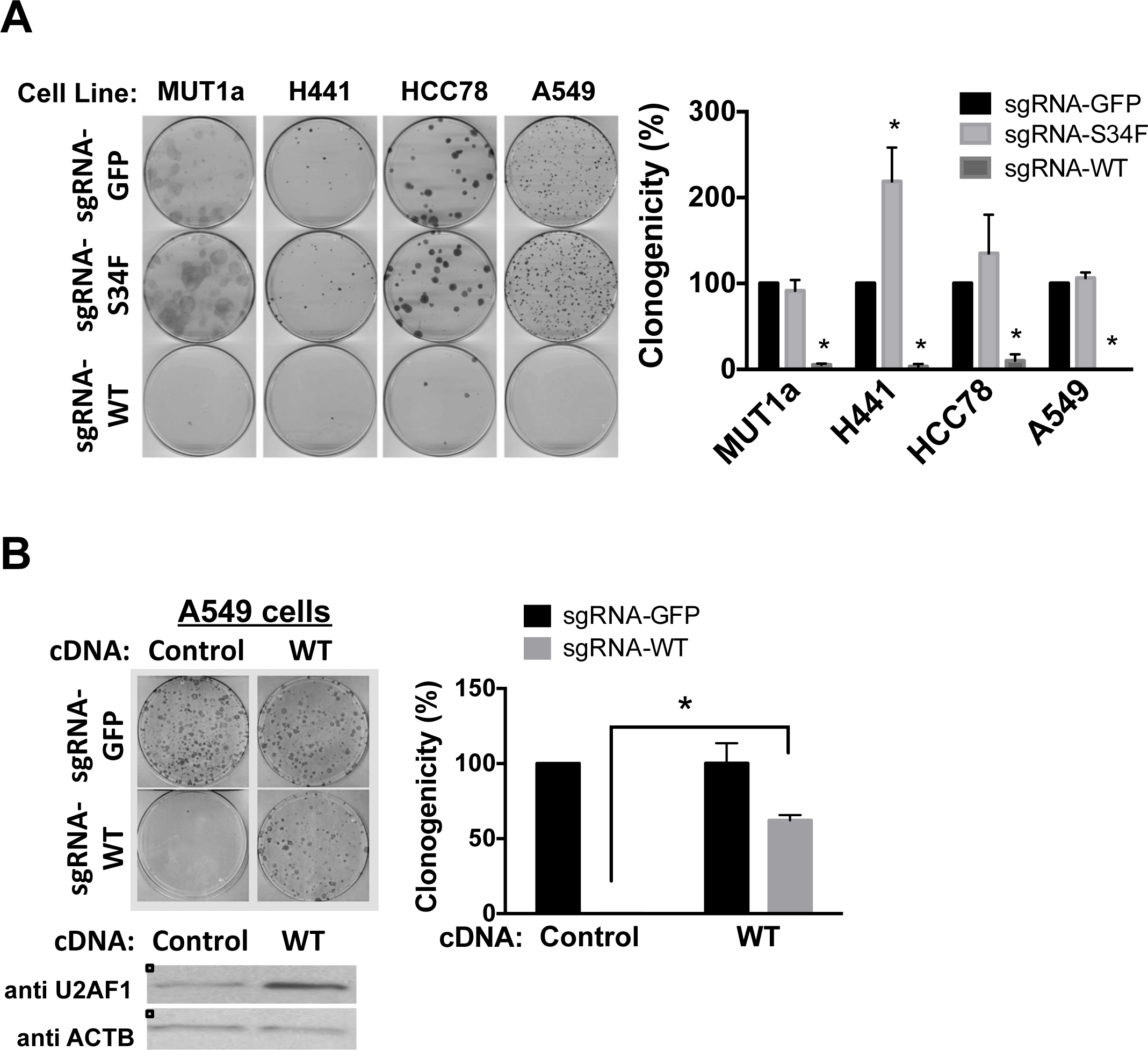
Wild-type but not mutant *U2AF1* is required for the clonogenic growth of the isogenic HBEC3kt cells and LUAD cell lines. (A) Clonogenic growth assays after selective disruption of the wild-type or mutant *U2AF1* allele. Left panel: The indicated cell lines were transduced with lentiviruses expressing Cas9 and sgRNA-GFP, sgRNA-S34F or sgRNA-WT, followed by clonogenic assays. Cell colonies were stained with methylene blue and counted three weeks later.Right panel: Quantification of the clonogenic assay. The results are shown as percent clonogenicity by setting the number of control cell colonies (cells transduced with Cas9 and sgRNA-GFP) as 100%. Asterisks represent statistical significant changes as compared to Cas9 and sgRNA-GFP-transduced condition in each cell line. Error bars represent s.e.m (n = 3). **(B)**. Rescue of growth inhibition by Cas9 and sgRNA-WT by 908 overexpressing a form of wild-type *U2AF1* cDNA that is not predicted to be the target for sgRNA-WT (See Supplemental Fig. S8 and Materials and Methods). Left panel: A549 cells were transduced with a control (DsRed-Express 2) or a wild-type *U2AF1* cDNA that is not predicted to be the target for sgRNA-WT. Increased expression of wild-type U2AF1 was confirmed by immunoblot (left bottom panel). These cells were subsequently transduced with Cas9 and either sgRNA-GFP or sgRNA-WT followed by clonogenic assays as in panel A (left upper panel).Right panel: Quantification as in Panel A. The asterisk represents a statistical significant change for the indicated comparison. Error bars represent s.e.m. (n = 3).

To examine the effect of *U2AF1*S34F on tumor growth *in vivo*, we derived H441 and HCC78 cells transduced with Cas9 and sgRNA-S34F or sgRNA-GFP as polyclonal pools or as clones (Supplemental Figs. S14, S15). The cell lines were verified to either carry or not carry the *U2AF1*S34F allele, and we confirmed that the Cas9 and sgRNA-S34F-transduced cells lost the S34F-associated splicing program (Supplemental Figs.S16, S17).

We inoculated these subclonal cell lines subcutaneously in nude mice and monitored xenograft tumor growth. The H441-derived cell lines, in which the *U2AF1*S34F allele was disrupted, were able to establish tumors *in vivo* at rates similar to those observed for tumor cells carrying the mutant allele (Supplemental Table S6). The HCC78-derived cell lines did not grow palpable tumors after xenografting within the observation period, so the requirement for *U2AF1*S34F *in vivo* could not be tested in that line. These experiments show that *U2AF1*S34F is dispensable for growth of these LUAD cell lines *in vivo*, a result consistent with the clonogenicity assays shown in Fig. 6.We conclude that *U2AF1*S34F appears to be neither sufficient nor necessary for lung cell transformation in these assays. In contrast, wild-type *U2AF1* is required for cell viability, consistent with the retention of a wild-type allele in human cancers carrying common *U2AF1* mutations.

## Discussion

In this report, we have examined the mechanistic and phenotypic consequences of the common *U2AF1*S34F mutation. Our data demonstrate that the S34F:WT ratio controls the quantitative consequences of the *U2AF1* mutation for splice site recognition, and suggest that differential RNA-binding affinities of the mutant and wild-type protein result in preferential recognition of specific 3’ splice sites. Moreover, our finding that wild-type *U2AF1* is required for cell survival irrespective of the presence or absence of *U2AF1*S34F explains the genetic observation that tumors always retain an expressed copy of the wild-type allele. Finally, we expect that the genetic models of *U2AF1*S34F that we derived from immortalized lung epithelial cells, as well as cell lines derived from lung adenocarcinomas with the mutation, will prove useful for future functional studies of this common mutation.

### The S34F:WT ratio controls S34F-associated splicing

U2AF1S34F is known to induce specific splicing alterations, but it is not known how these changes are regulated. We show that the ratio of S34F:WT U2AF1 gene products is a critical determinant of the magnitude of S34F-associated splicing. This conclusion was demonstrated in an isogenic lung epithelial cell line engineered to express *U2AF1*S34F from one of the two endogenous *U2AF1* loci, and was further supported by analyses of human LUAD transcriptomes carrying the *U2AF1*S34F allele. These results suggest that wild-type U2AF1 antagonizes the splicing program associated with the S34F mutation.

### Altered RNA binding affinities often explain S34F-associated splicing changes

We find that a major functional difference between purified S34F mutant and wild-type U2AF1 proteins resides in altered binding affinities for a subset of 3’ splice sites. The trends in the RNA sequence preferences of S34F-mutant U2AF1 are consistent with the preferred 3’ splice sites of S34F-affected transcripts (here and in [17–20, 28]), which we use as the signature of S34F-associated differential splicing (Fig. 1A).For oligonucleotides that showed significant changes in binding affinities, the S34F mutation typically reduced or enhanced respective binding of the U2AF1 splicing factor complexes to 3’ splice sites preceded by a −3U or −3C (Fig. 5 and [18]). In support of our findings for the relevant ternary complex of human U2AF1, U2AF2 and SF1 subunits, recent studies confirmed that the corresponding S34F mutation inhibited binding of the minimal fission yeast U2AF heterodimer to a “UAG” splice site RNA [21]. These S34F-altered RNA affinities are consistent with the location of the substituted amino acid in a zinc knuckle that may directly contact RNA [17]. Although the effects of the S34F mutation on binding 3’ splice sites preceded by −3A are variable, extrusion or alternative U2AF1 binding sites for disfavored nucleotides could occur in different sequence contexts by analogy with other RNA binding proteins [29, 30].

Several lines of evidence support the idea that U2AF1S34F is capable of initiating pre-mRNA splicing once it binds to RNA. Nuclear extracts of cells overexpressing mutant *U2AF1* can support *in vitro* splicing reactions more efficiently than nuclear extracts derived from cells overexpressing wild-type *U2AF1* for a minigene with a specific 3’ splice site sequence [17]. In addition, mutant U2AF1 can compensate for loss of the wild-type factor for the inclusion of some U2AF1-dependent cassette exons [31]. Our observations that the altered RNA-binding affinities correlate well with S34F-associated splicing for the majority of splice sites that we tested further suggest that mutant and wild-type U2AF1 are functionally equivalent for downstream steps of splicing (Fig. 5 and [18]).

### A working model of S34F-associated Splicing

In light of our findings and existing evidence from the literature, we propose a model wherein mutant U2AF1 drives differential splicing by favoring the recognition of one of two competing 3’ splice sites (Supplemental Fig. S18). This model is motivated by three key facts. First, alternative splicing, in contrast to constitutive splicing, necessarily results from implicit or explicit competition between splice sites. (For example, cassette exon recognition can involve competition between the 3’ splice sites of the cassette exon itself and a downstream constitutive exon.) Second, mutant and wild-type U2AF1 complexes have different binding specificities, largely due to their preferences for distinct nucleotides at the −3 position, that lead them to preferentially bind distinct 3’ splice sites. Third, mutant and wild-type U2AF1 are likely functionally equivalent once they bind to RNA. Therefore, altering the cellular ratio of mutant and wild-type U2AF1 changes the amount of total U2AF1 bound to a given 3’ splice site in a sequence-specific manner, thereby promoting or repressing that splice site relative to a competing 3’ splice site.

Our proposed model is not exclusive of other possible contributing effects, such as competitive binding of mutant and wild-type U2AF1 to a factor with low stoichiometry (Supplemental Fig. S18), or effects of U2AF1S34F on the kinetics of co-transcriptional splicing as suggested recently [32].Future studies are needed to resolve these possibilities.

### How are cells with the U2AF1 S34F mutation selected during oncogenesis?

*U2AF1*S34F is recurrently found in LUAD, other solid tumors, and myeloid disorders, suggesting that the mutant allele confers a physiological property that provides a selective advantage during neoplasia. A gene ontology (GO) analysis for genes that show S34F-associated alterative splicing in HBEC3kt-derived isogenic cells shows significant alterations in many biological processes such as mRNA processing,RNA splicing, G2/M transition of mitotic cell cycle, double-strand break repair and organelle assembly (FDR-adjusted p-values < 0.001). However, we did not observe signs of neoplastic transformation or changes in cell proliferation after introducing *U2AF1*S34F into the endogenous *U2AF1* locus in HBEC3kt cells (Supplemental Fig.S13). Moreover, targeted inactivation of *U2AF1*S34F in LUAD cell lines does not diminish, and in one case even increases, clonogenic growth in culture (Fig. 6).

The lack of a testable cellular phenotype has been a major hindrance to understanding the functional significance of mutant *U2AF1* in carcinogenesis. Cell proliferation is only one of the many hallmarks of cancer, so careful examination of other cell properties in the isogenic cells may be needed to establish the presumptive role of *U2AF1*S34F in carcinogenesis.

More recently, after completion of our study [33], Park *et al* reported that tumorigenic cells emerge after *U2AF1*S34F is ectopically produced in Ba/F3 pro-B cells or in an immortalized line of small airway epithelial cells [34]. The authors attributed the transformation events by mutant U2AF1 to the consequences of altered 3’ processing of mRNA’s. In particular, they observed an increase in the length of the 3’ untranslated region of *ATG7* mRNA and a reduction in the amount of ATG7 protein, proposing that the anticipated defect in autophagy predisposes cells to mutations, some of which are transforming. This “hit-and-run” type of mechanism is consistent with our observation and theirs that mutant U2AF1 is dispensable for maintenance of the transformed phenotype in LUAD cell lines (Figure 6) and in their cell lines [34]. Their observations do not, moreover, exclude a role for S34F-associated splicing in the oncogenic mechanism.

### Can cancer cells carrying the S34F mutation be targeted therapeutically?

We have shown that the wild-type *U2AF1* allele is absolutely required for the growth of lung epithelial and LUAD cells that carry a mutant *U2AF1* allele (Fig. 6). This result indicates that mutant U2AF1 cannot complement a deficiency of wild-type U2AF1; it may also explain why tumors homozygous for the *U2AF1*S34F mutation have not been observed, although the number of tumors found to have even a heterozygous mutation is still relatively small, so the analysis may not be adequately powered. Still, the frequent occurrence of a low ratio of S34F:WT *U2AF1* mRNA, accompanied by increased copy number of wild-type *U2AF1* alleles in lung adenocarcinoma cell lines and possibly LUADs, suggests that there may be selection for a lower ratio of S34F:WT in addition to the likely selection, perhaps at an earlier stage of tumorigenesis, for the S34F mutant.

These results are consistent with a study of mutations affecting the splicing factor SF3B1, which reported that cancer cells harboring recurrent *SF3B1* mutations also depend on wild-type SF3B1 for growth [35]. Finally, a recent study similarly found that a wild-type copy of *SRSF2* is required for leukemic cell survival, and that *SRSF2* mutations generated a therapeutic index for treatment with a small molecule that inhibits 3’ splice site recognition [36]. Together, our results and these recent studies provide a genetic rationale for targeting wild-type splicing factors (or the splicing machinery more generally) in cancers harboring spliceosomal mutations.

## Materials and Methods

### Cell culture, reagents, and assays

The HBEC3kt cells (a gift from Dr. John Minna, UT Southwestern Medical Center), H441, A549 (ATCC) and HCC78 cells (DMSZ) were cultured as previously described [22] or according to vendors’ instructions. The primary antibodies for immunoblots are: rabbit anti U2AF1 (1:5000, # NBP1-19121, Novus), rabbit anti GFP(1:5000, #A-11122, Invitrogen), mouse anti ACTB (1:5000, Clone 8H10D10, Cell Signaling). Lentiviruses were produced and titered in HEK293T cells as previously described [37]. An MOI (multiplicity of infection) of 1 – 5 were used for all assays.

Total RNA was extracted and reverse transcribed as previously described [38].Splicing alterations were measured by quantitative PCR on reverse-transcribed cDNA (RT-qPCR), using isoform-specific primers (Supplemental Table S7). These primers were designed following a previously described method [39]. The PCR efficiency and specificity of each primer set were determined before they were used for measuring splicing changes (See Supplemental Materials and Methods for details).

Clonogenic assay was performed by infecting cells with lentiviruses two days before seeding them into 100 mm dishes (1000 live cells per dish) to grow colonies. Growth media were supplemented with puromycin (1 µlg/ml) for selecting infected cells and were changed once a week for up to three weeks. Cell colonies were stained with 0.03% methylene blue (in 20% methanol) for 5 min. Clonogenicity is defined as colony 598 numbers formed as a percent of those in control cells.

Details of all DNA constructs used in the study and the genome editing approaches for creating the *U2AF1*S34F allele and allelic-specific disruption of *U2AF1* are described in the Supplemental Materials and Methods.

### mRNA sequencing and analysis

High throughput mRNA sequencing (RNA-seq) was conducted in the Sequencing Facility of the National Cancer Institute. RNA quality was assessed by 2100 Bioanalyzer (Agilent). Total RNA with good integrity values (RIN > 9.0) was used for poly A selection and library preparation using the Illumina TruSeq RNA library prep kit. Two or four samples were pooled per lane and ran on the HiSeq2000 or HiSeq2500 instrument using TruSeq V3 chemistry. All samples were sequenced to the depth of 100 million pair-end 101 bp reads per sample.

Splicing analysis of RNA-seq data from the TCGA LUAD cohort as well as engineered HBEC3kt, H441, and HCC78 cell lines was performed as previously described [17]. A brief description of the method was provided in the Supplemental Materials and Methods.

### Purification of U2AF1 protein complexes and RNA affinity measurement

Purification of U2AF1 complexes, as illustrated in Fig.5, was explained in Supplemental Materials and Methods. Sequences of synthetic 5’-labeled fluorescein RNAs (GE Healthcare Dharmacon) and binding curves are given in the Supplementary Fig. S12. Apparent equilibrium affinity constant of the purified U2AF1 complexes with RNA was measured based on changes in fluorescence anisotropy as previously described [18]. A brief description of this method was provided in the Supplemental Materials and Methods.

### Statistics

All experiments were independently performed at least three times unless otherwise stated. Statistical significance was determined by two-tailed Student’s *t* test or otherwise stated. In all analyses, *p* values ≤ 0.05 are considered statistically significant.

## Acknowledgements

We thank Ms. Jackie Idol, Ursula Harper, Danielle Miller-O’Mard and the transgenic mouse core at NHGRI for technical assistance, Dr. Heidi Dvinge (FHCRC) for help with the LUAD data analysis, Dr. Matthew J. Walter (Wash.U.) for sharing the *CEP164* and *FMR1* splice site sequences prior to publication. We thank members of the Varmus lab, Drs. Janine Ilagan (FHCRC) and Paul Liu (NHGRI) for helpful discussions during the course of the study. HV was supported by the Intramural Program at the National Institutes of Health and is now supported by the Meyer Cancer Center at Weill Cornell Medicine. RKB is supported by the Edward P. Evans Foundation, Ellison Medical Foundation (AG-NS-1030-13), NIH/NHLBI (R01 HL128239), and NIH/NIDDK (R01 DK103854). CLK is supported by the Edward P. Evans Foundation and NIH/NIGMS (R01 GM070503). The results published here are in part based upon data generated by the TCGA Research Network: http://cancergenome.nih.gov/.

